# Hawkmoth pheromone transduction involves G protein-dependent phospholipase Cβ signaling

**DOI:** 10.1101/2024.08.29.610295

**Authors:** Anna C. Schneider, Katrin Schröder, Yajun Chang, Andreas Nolte, Petra Gawalek, Monika Stengl

## Abstract

1

Evolutionary pressures adapted insect chemosensation to the respective insect’s physiological needs and tasks in their ecological niches. Solitary nocturnal moths rely on their acute olfactory sense to find mates at night. Pheromones are detected with maximized sensitivity and high temporal resolution through mechanisms that are mostly unknown. While the inverse topology of insect olfactory receptors and heteromerization with the coreceptor Orco suggest ionotropic transduction via odorant-gated receptor-ion channel complexes, contradictory data propose amplifying G protein-coupled transduction. Here, we used *in vivo* tip-recordings of pheromone-sensitive sensilla of male *Manduca sexta* hawkmoths at specific times of day (rest vs. activity). Since the olfactory receptor neurons distinguish signal parameters in three consecutive temporal windows of their pheromone response (phasic; tonic; late, long-lasting), respective response parameters were analyzed separately. Disruption of G protein-coupled transduction and block of phospholipase C decreased and slowed the phasic response component during the activity phase of hawkmoths without affecting any other component of the response during activity and rest. A more targeted disruption of G_α_ subunits by blocking G_αo_ or sustained activation of G_αs_ using bacterial toxins affected the phasic pheromone response, while toxins targeting G_αq_ and G_α12/13_ were ineffective. Consistent with these data, the expression of phospholipase Cβ4 depended on zeitgeber time, which indicates circadian clock-modulated metabotropic pheromone transduction cascades that maximize sensitivity and temporal resolution of pheromone transduction during the hawkmoth’s activity phase. Thus, discrepancies in the literature on insect olfaction may be resolved by considering circadian timing and the distinct odor response components.

**2 Significance statement:** Insect chemosensory transduction is typically thought to be ionotropic, but data from different insect species suggests that metabotropic olfactory signaling may occur, either alongside or instead of ionotropic mechanisms. Nocturnal moths, known for their extraordinarily sensitive pheromone-detecting olfactory receptor neurons, likely use metabotropic signal amplification. To overcome limitations of previous *in vitro* studies, we conducted tip-recordings of pheromone-sensitive sensilla in healthy hawkmoths at specific zeitgeber times. Disrupting G protein signaling and phospholipase Cβ reduced sensitivity and altered response kinetics, revealing strict temporal control of transduction. Thus, contradictory findings in insect olfaction may be reconciled by considering diverse evolutionary pressures for distinct chemosensory signals in different species, zeitgeber time, and disparate odor response parameters.

## 3 Introduction

Both male and female nocturnal *Manduca sexta* hawkmoths express daily rhythms in their mating behavior, which are synchronized via species-specific pheromones that are released exclusively at night. Superimposed on this slow, nightly rhythm of pheromone release, female moths emit their sex pheromone blend in fast pulses via oscillatory exposure of their abdominal pheromone glands (Riffell et al., 2008; Sasaki and Riddiford, 1984; Schendzielorz et al., 2015; Tumlinson et al., 1989). The males detect intermittent changes of the pheromone blend with long trichoid sensilla on their antenna, each of which is innervated by two olfactory receptor neurons (ORNs). One of the two ORNs specifically detects bombykal (BAL), the main sex pheromone component (Altner and Prillinger, 1980; Kaissling et al., 1989; Keil, 1989; Keil and Steinbrecht, 1984; Sanes and Hildebrand, 1976).

Pheromone-sensitive ORNs of moths are outstandingly sensitive with a dynamic range that covers approximately 7 log units of pheromone concentrations. They are flux detectors, tuned to resolve fast changes in pheromone concentration (Baker and Vogt, 1988; Kaissling, 1998; Marion-Poll and Tobin, 1992). Their sensitivity and electrical response kinetics depend on stimulus strength and duration. ORN properties are controlled by their endogenous circadian clocks, which results in maximum sensitivity and fastest pheromone pulse tracking at night, during the hawkmoth’s activity phase (Dolzer et al., 2003; Flecke et al., 2010; Flecke and Stengl, 2009; Schuckel et al., 2007). The molecular mechanisms, which are responsible for their exceptional sensitivity and which can be adjusted by zeitgeber time (ZT) and stimulus intensity, are poorly understood and contradictory findings for insect pheromone/odor transduction cascades have been reported, even in the same species and for the same odorants (Nakagawa and Vosshall, 2009; Stengl, 2017, 2010; Stengl and Funk, 2013; Wicher and Miazzi, 2021).

Insect olfactory receptors (ORs) are 7 transmembrane (7TM) molecules with inverse topology and not related to the G protein-coupled 7TM ORs of vertebrates (Benton et al., 2006; Clyne et al., 1999; Gao and Chess, 1999; Krieger et al., 2003, 2002; Lundin et al., 2007; Vosshall et al., 1999). In *Drosophila melanogaster*, ORs heteromerize with a distantly related but conserved inverse 7TM molecule, Or83b, the olfactory receptor coreceptor (Orco) (Vosshall et al., 1999; Vosshall and Hansson, 2011). While for *D. melanogaster* both an ionotropic and mixed ionotropic and metabotropic odor/pheromone transduction cascade has been suggested (Sato et al., 2008; Wicher et al., 2008), no evidence for OR-Orco-dependent ionotropic pheromone transduction was found in *M. sexta* (Nolte et al., 2016, 2013). In *D. melanogaster*, OR-Orco complexes constitute odorant-gated, non-specific cation channels with slow gating kinetics and a reversal potential around 0 mV (Sato et al., 2008; Wicher et al., 2008). In contrast, in *M. sexta*, the first ion channel that activates after pheromone application is a transient Ca^2+^ channel with rapid kinetics that opens and closes in less than 100 ms. The resulting pheromone-dependent current resembles the inositol trisphosphate (IP_3_)- or diacyl glycerol (DAG)-dependent currents that are reminiscent of TRP ion channel family-dependent currents (Dolzer et al., 2021; Gawalek and Stengl, 2018; Stengl, 1994, 1993; Stengl et al., 1992). Thus, in hawkmoths, pheromone transduction was suggested to employ a metabotropic G protein-coupled cascade that activates phospholipase Cβ (PLCβ), which allows strong signal amplification and a flexible, adjustable dynamic range (Stengl, 2010).

In contrast to most of the previous research on insect odor/pheromone transduction cascades, here, we used *in vivo* experiments in healthy *M. sexta* hawkmoths to challenge the hypothesis of metabotropic PLCβ -dependent pheromone transduction. We combined tip-recordings of pheromone-sensitive long trichoid sensilla with pharmacology in search for circadian modifications of pheromone transduction that are not directly mediated by light levels. We compared the ORN responses to BAL stimulation at the end of the hawkmoth’s activity phase at ZT 1-3 to the response at the resting phase at ZT 9-11. The BAL-elicited spiking response consists of three consecutive components with distinct firing patterns and durations. Each of the components carries different information of the pheromone signal (Dolzer et al., 2003; Nolte et al., 2016, 2013). Changes in pheromone concentration are encoded only by the latency and spike frequency of the first component, the phasic response (about 10 spikes) (Dolzer et al., 2003). Thus, a comprehensive data analysis of the distinct components of the pheromone/odor response is necessary because, while being conserved across insects, their distinct kinetics indicate different underlying mechanisms of insect olfaction (Dolzer et al., 2003; Kaissling, 1987, 1986; Nolte et al., 2016, 2013; Stengl, 2017; Stengl and Funk, 2013). Therefore, we carefully distinguished response parameters, both in terms of response component and ZT.

## 4 Methods

### 4.1 Animals

For all experiments, we used two- to three-day old adult male *Manduca sexta* (Lepidoptera, Sphingidae) hawkmoths from the breeding and rearing colonies at the University of Kassel, Germany. Animals were raised on an artificial diet and kept under long-day conditions (L:D; 17:7 h) at 24-26 °C and relative humidity of 40-60 %. After pupation, males and females were separated and males were housed isolated from pheromones. Animals were fed and raised as described in Gawalek and Stengl (2018).

### 4.2 Solutions

Salts for ringer solutions were purchased from Merck. Bacterial toxins, GDP-β-S, U73122 and U73343 were purchased from Sigma-Aldrich. All other chemicals were obtained from Sigma, unless stated otherwise.

Ringer solutions were prepared as described previously (Gawalek and Stengl, 2018). In brief, sensillum lymph ringer (SLR) contained (in mM): 172 KCl, 3 MgCl_2_, 1 CaCl_2_, 25 NaCl, 10 HEPES, 22.5 glucose, pH 6.5 and adjusted to 475 mOsmol/kg with glucose. Hemolymph ringer (HLR) contained (in mM): 6.4 KCl, 12MgCL_2_, 1 CaCl_2_, 12 NaCl, 10 HEPES, 340 glucose, pH 6.5 and adjusted to 450 mOsmol/kg with glucose.

Synthetic bombykal (BAL; E,Z-10,12-hexadecadienal; gift from J. H. Tumlison, Center for Medical, Agricultural and Veterinary Entomology) was prepared as stock solution of 100 µg/ml in n-hexane and stored at -80 °C until use.

As general inhibitor of G proteins, we used 10^-5^ M GDP-β-S (10^-2^ M stock solution dissolved in DMSO, Sigma-Aldrich) in SLR. For further discrimination between G protein subunits, we used the following bacterial toxins: pertussis toxin (pTox, 500ng/500µl solution in SLR, stored at 4 °C until use) is a specific inhibitor of G_αo_ in *D. melanogaster* (Hopkins et al., 1988) and blocks IP_3_ production in cockroach (Boekhoff et al., 1990); cholera toxin (cTox, 5 µg/500µl solution in SLR, stored at 4 °C until use) activates G_αs_ in insects (Deng et al., 2011), and *Pasteurella multocida* toxin (pmTox, 500 ng/500 µl solution dissolved in SLR, stored at -20 °C until use) activates G_αq_ and G_αi/o_ (Surguy et al., 2014). For disruption of phospholipase C (PLC) signaling, we used the antagonist U73122 and its respective non-functional analog U73343 (both prepared as 10^-2^ M stock solution dissolved in DMSO and stored at -80 °C until use), diluted to a final a concentration of 10^-5^ M in SLR.

### 4.3 Electrophysiology

To investigate the molecular mechanisms of the pheromone signal transduction cascade, we performed long-term (hours to days) tip-recordings of the ORNs of single antennal long trichoid sensilla (Kaissling, 1995) at room temperature (20-23 °C). Animals were fixed in a custom-made holder with tape, and the antenna was immobilized with dental wax (Boxing Wax, Kerr) near the base. To record from ORNs, the recording electrode was slipped over the clipped tip of one long trichoid sensillum and the reference electrode was inserted into the clipped antenna close to the annulus where the recording electrode was located. Electrodes were moved with manual micromanipulators (MMJ, Märzhäuser Wetzlar; Leitz, Leica Microsystems). Electrodes were pulled from thin-walled borosilicate glass capillaries (OD 1.5 mm, ID: 1.17 mm, Havard Apparatus) with a tip opening that both sealed well around the sensillum and fit the inside of the antenna (∼1 µm). The cut antenna was sealed with electrode gel (GE Medical Systems Information Technology) to prevent desiccation. The recording electrode was filled with SLR, the reference electrode with HLR. ORN activity was amplified 200-fold (custom build amplifier with 10^12^ Ω input impedance, or BA-03X, npi, or ELC-01MX, npi), low pass filtered with a cut-off frequency of 1.3 kHz, digitized at 20 kHz with a Digidata (1200, or 1550A or 1550B, Molecular Devices), and saved for offline analysis with pClamp 8 or 10 (Molecular Devices). We used pClamp to record the signal as high-pass filtered (150 Hz cut-off frequency) AC signal in addition to the DC signal.

For BAL stimulation, 10 µl stock solution were loaded on filter paper (∼1 cm^2^), resulting in 100 ng BAL per filter paper; air streams for stimulation were set up according to previously described protocols (Gawalek and Stengl, 2018; Nolte et al., 2013). BAL stimulation consisted of periodic application of a 50 ms pulse at an interstimulus interval of 5 min to avoid desensitization and habituation. Recordings with SLR in the recording electrode served as controls, either with or without the addition of 0.1% DMSO, as specified. The addition of 0.1% DMSO to the SLR did not alter the recorded BAL response. Activators, inhibitors, or toxins were added to the SLR of the recording electrode.

To search for zeitgeber time (ZT) dependent effects, we recorded during the animal’s late activity phase (ZT 1-3) and in the middle of its resting phase (ZT 9-11). In experiments involving G protein or PLC signaling disruption, we recorded from different animals at the different ZTs and different interference conditions. Recording and periodic stimulation usually lasted for 120 min, with a few exceptions in the control recordings, which only lasted for about 70 min. In the toxin experiments, we did paired recordings of the same animal at the same ZTs on two consecutive days. For these experiments, we always filled the recording electrode with toxin SLR, obtained the control recordings on day 1 immediately after attaching the electrode but before the toxins had any effect, and recorded on the same ZT on day 2 without removing the electrodes in between. Here, we stimulated three times in the control recording and six times in the toxin recording.

### 4.4 Tissue collection, RNA extraction, and cDNA synthesis

We collected antennal tissue from isolated cultures of 2-day old males. At each ZT1, ZT9, and ZT17, we collected three samples with 8 antennae per sample. All samples were immediately frozen in liquid nitrogen and stored at -80 ℃ for RNA extraction. Total RNA was isolated using TRIzol® Reagent (Invitrogen) according to the manufacturer’s protocol. The quality of the RNA was assessed by gel electrophoresis. The concentration and purity of RNA were detected using NanoDrop ND-1000 spectrophotometer (Thermo Fisher Scientific). OD_260/280_ ratios of all samples were 1.8 - 2.0. The first-strand complementary DNA (cDNA) was synthesized with 1 μg of total RNA for each sample using the PrimeScript™ RT reagent Kit with gDNA Erase (Takara Bio Inc.) according to the manufacturer’s instructions and stored at -20 ℃ until use.

### 4.5 Phylogenetic analysis of *PLCβ* from *M. sexta* and other species

Since the *PLCβ* gene is not annotated clearly in the genome of *M. sexta*, we utilized the coding sequences (cds) of the *PLCβ* genes from *D. melanogaster* and *Bombyx mori*. We identified two highly similar genes in *M. sexta* as candidate *PLCβ1* (Gene ID: LOC115440592) and *PLCβ4* (*norpA*; Gene ID: LOC115451385) with BLAST analysis in the NCBI database. We downloaded the cds of these candidate genes, and the *PLC-β* genes of other insects, *Homo sapiens* and *Mus musculus*.

We translated the cds into protein sequences and performed sequence alignments using MEGA X (Kumar et al., 2018). The phylogenetic tree was constructed using the maximum likelihood method and Tamura-Nei model (Tamura and Nei, 1993), and node support was evaluated with 1000 bootstrap replicates.

### 4.6 Real-time quantitative polymerase chain reaction (qPCR)

qPCR was conducted to determine the expression of target genes (Table 1) at ZT1, ZT9, and ZT17 of hawkmoth antennae with TB Green® Premix Ex Taq™ II (Tli RNase H Plus, Takara Bio Inc.). Primers for each target gene were designed using the Primer-BLAST online program of NCBI (Table 1). *Ribosomal Protein S13* (*RPS13*) (Fenske et al., 2018) and *Glyceraldehyde-3-Phosphate Dehydrogenase* (*G3PDH*) (Adamo et al., 2016; Mészáros and Morton, 1996) were chosen as the reference genes to normalize the relative expression levels of each target gene. Agar gel electrophoresis, sequencing of PCR fragments and subsequent qPCR showed that the expression levels of the reference genes in the antennae were stable at ZT1, ZT9, and ZT17. In this study, all primers were used to amplify the target genes via PCR with cDNA. Following amplification, the PCR products were resolved by gel electrophoresis. Target bands were excised and purified using NucleoSpin Gel and PCR Clean-up Kit (Macherey-Nagel GmbH & Co. KG). The purified DNA was subsequently sequenced using the Sanger method to verify the specificity of the amplified fragments.

**Table 1:**
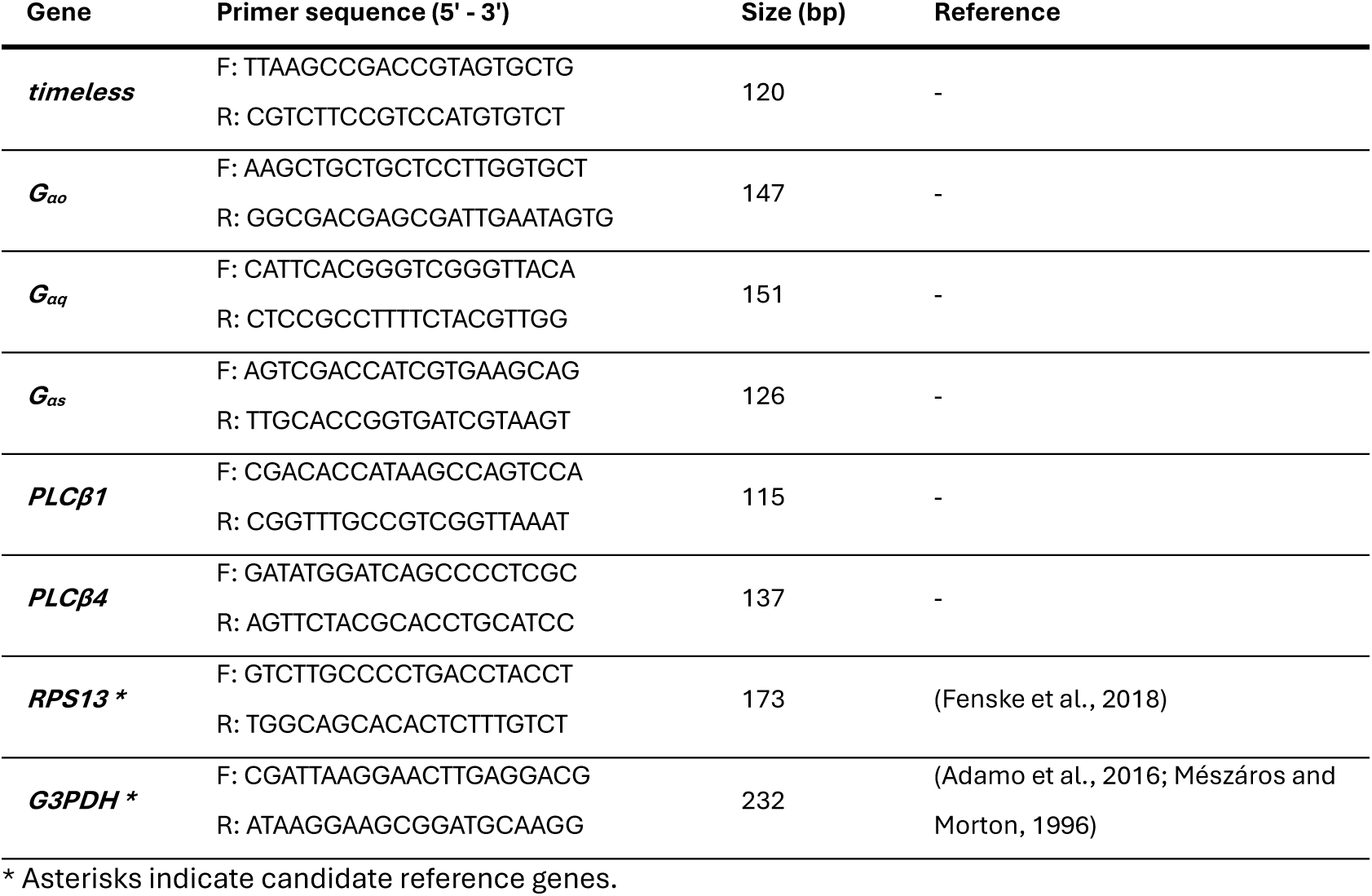
Forward and reverse primer sequences for target and reference genes.

The qPCR reaction program was: 95 °C for 30 s, followed by 40 cycles of 95 °C for 5 s, 60 °C for 30 s. For each gene and biological repeat three technical replicates. The amplification efficiency of all primers was 90 % - 110 %. The relative expression levels of these target genes were calculated according to the 2^-ΔΔCt^ method (Livak and Schmittgen, 2001).

### 4.7 Data analysis

For evaluation of pheromone responses, both the DC and high-pass filtered signal (AC) were examined. Spikes were detected by a threshold search on the AC trace. During the first second after BAL stimulation, the following pheromone response parameters were evaluated (Figure 1A): latency of the first spike relative to the beginning of the DC pheromone response, phasic spike frequency as the average instantaneous frequency of the first six spikes (*F*_6AP_), number of spikes in a 1 s window (general G protein and PLC disruption experiments) or 100 ms window (toxin experiments) (#APs (early)), and sensillum potential amplitude (SPA). The SPA was measured as the maximal deflection of the transepithelial potential following the BAL stimulation. Furthermore, for analysis of response kinetics, all APs within the first second after BAL stimulation were counted and binned in 10 ms bins. For the late long-lasting pheromone response (LLPR) that persists over several minutes after pheromone stimulation, we counted the number of spikes in the 295 s before the next BAL stimulus Figure 1B.

**Figure 1:**
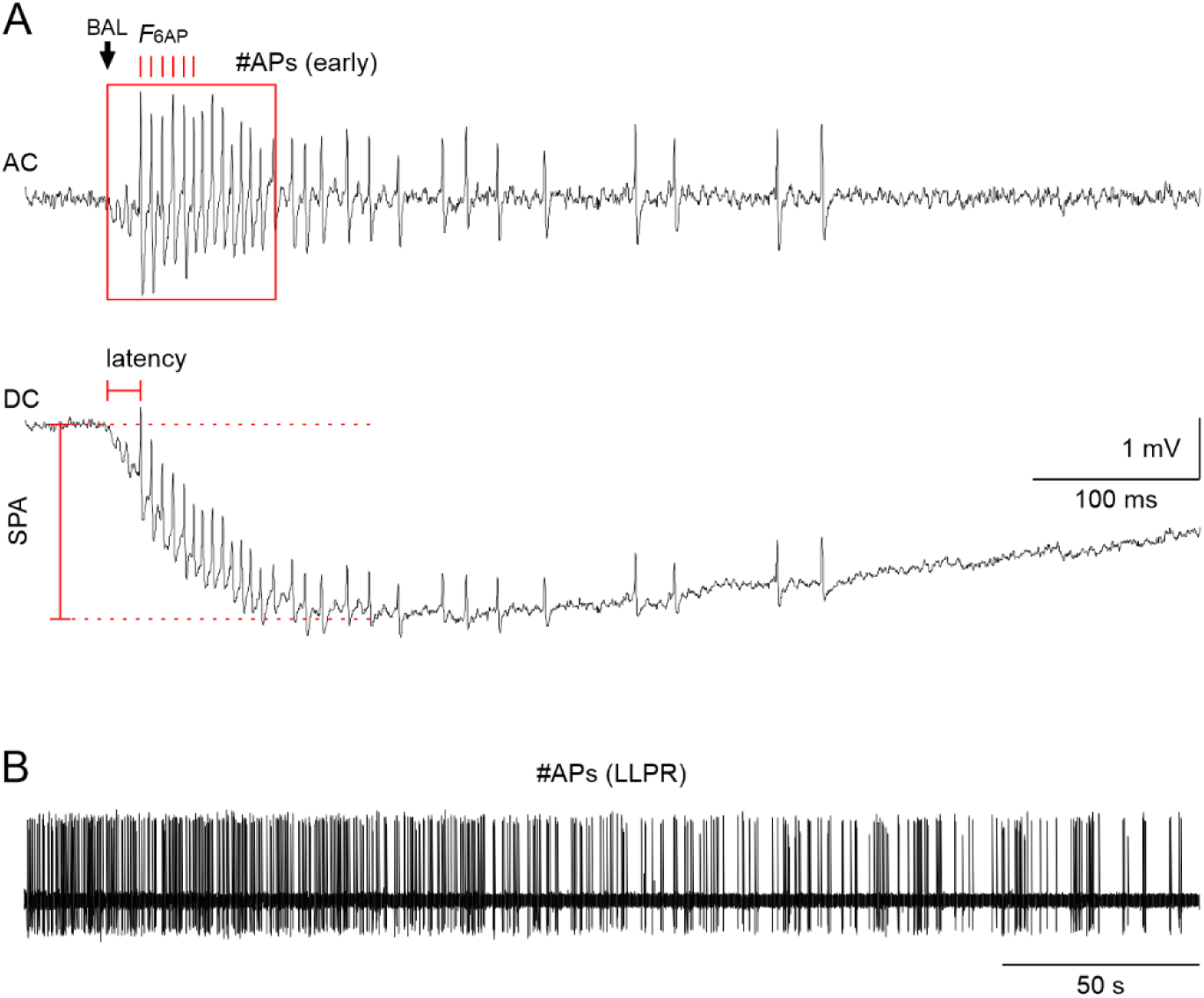
Quantification of the three phases of the pheromone response of hawkmoth long trichoid sensilla. The bombykal (BAL)-elicited spiking response consists of three consecutive spiking patterns (3 components) with distinct kinetics but variable duration:1) a phasic, high-frequency spiking response that lasts less than 100 ms (**A**), 2) a tonic spiking response with lower and more variable spiking frequency that lasts more than 100 ms (**A**), and 3) a late long-lasting spiking response (LLPR) that lasts seconds to minutes (**B**) (Dolzer et al., 2003; Nolte et al., 2016). **(A)** High-pass filtered (AC, top trace) and unfiltered (DC, bottom trace) recording of the sensillum potential with action potentials (APs) of the phasic-tonic ORN response to BAL. BAL stimulus: 50 ms, 10 µl of 0.1 mg/ml on 1 cm^2^ filter paper; arrow: start of the BAL response. For the phasic response we calculated the average instantaneous AP frequency of the first six APs (*F*_6AP_, red ticks, top trace) of the BAL response. A combination of both phasic and part of the tonic response was evaluated as the number of APs in a 100 ms window starting at the onset of the BAL response (#APs (early), red box, top trace). Latency (red horizontal marking, bottom trace) is the time from the beginning of the BAL response to the first AP. Sensillum potential amplitude (SPA) is the amplitude from the baseline voltage before BAL stimulation to the negative peak (red vertical marking, bottom trace) **(B)** High-pass filtered recording of the LLPR. The LLPR was evaluated as the number of APs (#APs (LLPR)) in the 295 s before the next BAL stimulus, excluding the first 5 s after the BAL stimulus.

In experiments involving G protein or PLC signaling disruption, the time course of the responses to repeated stimulations varied across animals, presumably due to individual differences in the reaction to DMSO application *in vivo* or differences between individual sensilla. First effects were usually observed within 10-30 min after placement of the electrodes but due to the *in vivo* recording method the final concentration of the chemicals in the sensillum lymph was unknown. Therefore, we used linear fits to describe the change of each response parameter over the 120 min of stimulation. The slopes of the fits were used for quantification (Figure 2A). Slopes that were not significantly different from zero (no significant change of the response over time, t-test) were set to zero.

**Figure 2:**
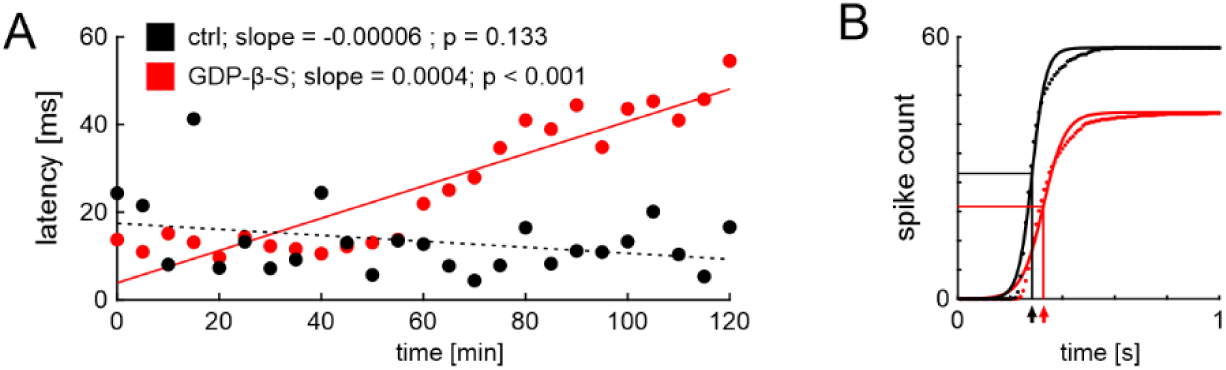
Illustration of the linear fit analysis of pheromone responses. **(A)** Bombykal stimuli were applied every 5 min in the 120 min long tip-recordings of hawkmoth trichoid sensilla. For each animal and experimental condition (here: one control (black) and one GDP-ϐ-S (red) animal) we generated a linear regression model for time and respective BAL response parameters (here: latency to the first action potential after the onset of the BAL response) and used a t-test to determine whether the slope of the fit significantly differed from zero (solid line) or not (dashed line). **(B)** To quantify the response kinetics to BAL stimulation, we binned the data of the first second after the BAL stimulus in 10 ms bins across all BAL stimulations for each animal. Each resulting cumulative histogram (dotted line) was fitted with a sigmoidal function (solid line) for quantification that yielded three fit parameters: maximum spike number (AP_max_), time of the midpoint of the sigmoid (t_1/2_; indicated by arrows at the x-axis), and slope of the midpoint (k). Depicted are examples of two animals.

To quantify the changes of the kinetics of the spiking response to BAL stimulation we created cumulative histograms of the binned data and fitted those histograms with the sigmoidal function

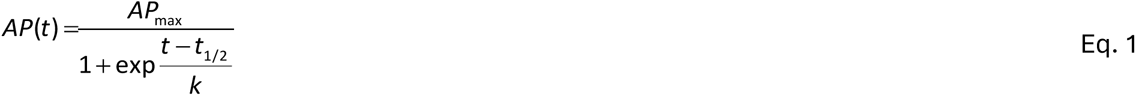

where *AP_max_* is the cumulative number of spikes within 1 s after stimulation, *t_1/2_* is the time of the midpoint, and *k* the slope factor (Figure 2B). The fit parameters were used for quantification.

### 4.8 Statistics

Data were tested for normal distribution with the Shapiro-Wilk test and for equal variance with Levene’s test. We used 1-way ANOVA or repeated-measures (RM) ANOVA for normally distributed data with equal variance as indicated. If data were not normally distributed, we log-transformed the data or used ANOVA on ranks if transformation did not result in a normal distribution. For pairwise comparisons, we used Tukey’s post-hoc test for normally distributed and Dunn’s post-hoc test for non-normally distributed data. All tests were done in SigmaPlot (version 12, Systat Software).

## 5 Results

The response of antennal pheromone-sensitive trichoid sensilla to BAL stimulation was recorded *in vivo* in restrained, intact male hawkmoths at two zeitgeber times: at the end of the activity phase (ZT 1-3) and in the middle of the resting phase (ZT 9-11). Pharmacology was used to determine whether hawkmoth ORNs comprise G protein-coupled pheromone receptors that activate PLCϐ, and whether this pathway is under the control of an ORN-based endogenous circadian clock to maximize sensitivity and temporal resolution of pheromone detection during the activity phase.

The pheromone response of ORNs consists of three components. The first component is fast and phasic, and comprises less than 10 spikes (APs) which encode BAL concentration changes (Dolzer et al., 2003). It is characterized by regular, high-frequency spiking in a time window of ≤ 100 ms. The second component is tonic spiking at a lower and more variable frequency, which does not encode pheromone concentration changes but encodes signal duration within a limited range. It typically lasts several hundreds of milliseconds. This is followed by a pause and then the third component, the spiking of the late long-lasting pheromone response (LLPR), which can last seconds to minutes. The LLPR encodes neither quantity nor duration of the pheromone signal. It is distinct from the spontaneous activity before pheromone application and is hypothesized to encode a memory trace of the previous pheromone signal (Stengl, 2010).

### 5.1 General inhibition of G proteins reduced the spike frequency and increased response latency of the phasic component of the BAL response during the activity phase but not during rest

To search for G protein coupling in the pheromone transduction cascade, 10 µM GDP-ϐ-S, an indiscriminative inhibitor of G proteins, were infused via the tip-recording electrode into the pheromone-sensitive trichoid sensillum. The ORNs’ neural responses to BAL stimulation were recorded with a non-adapting, periodic stimulation protocol at two different ZTs in control and with GDP-β-S.

During the activity phase, blocking of G protein-coupled processes reduced the frequency of the phasic response component (*F*_6AP_), while the latency increased (Figure 3A, B, D; ANOVA: Table 2; raw data: Extended Data Figure 3-1). In contrast, GDP-β-S had no significant effect on *F*_6AP_ or latency during the resting phase (Figure 3B-D; ANOVA: Table 2; raw data: Extended Data Figure 3-1). However, response variability across animals increased in GDP-β-S at both ZTs over time, as indicated by the large increase in standard deviation (Figure 3B, C). In addition, the kinetics of the response, i.e., rise and fall times to the maximum spike rate, were diminished with GDP-β-S during the activity phase but not at rest, as indicated by the larger slope factor *k* of the sigmoidal fits (Figure 4; ANOVA: Table 3; raw data: Extended Data Figure 4-1). GDP-ϐ-S application did not have significant effects on any of the other response parameters at either ZT.

**Figure 3:**
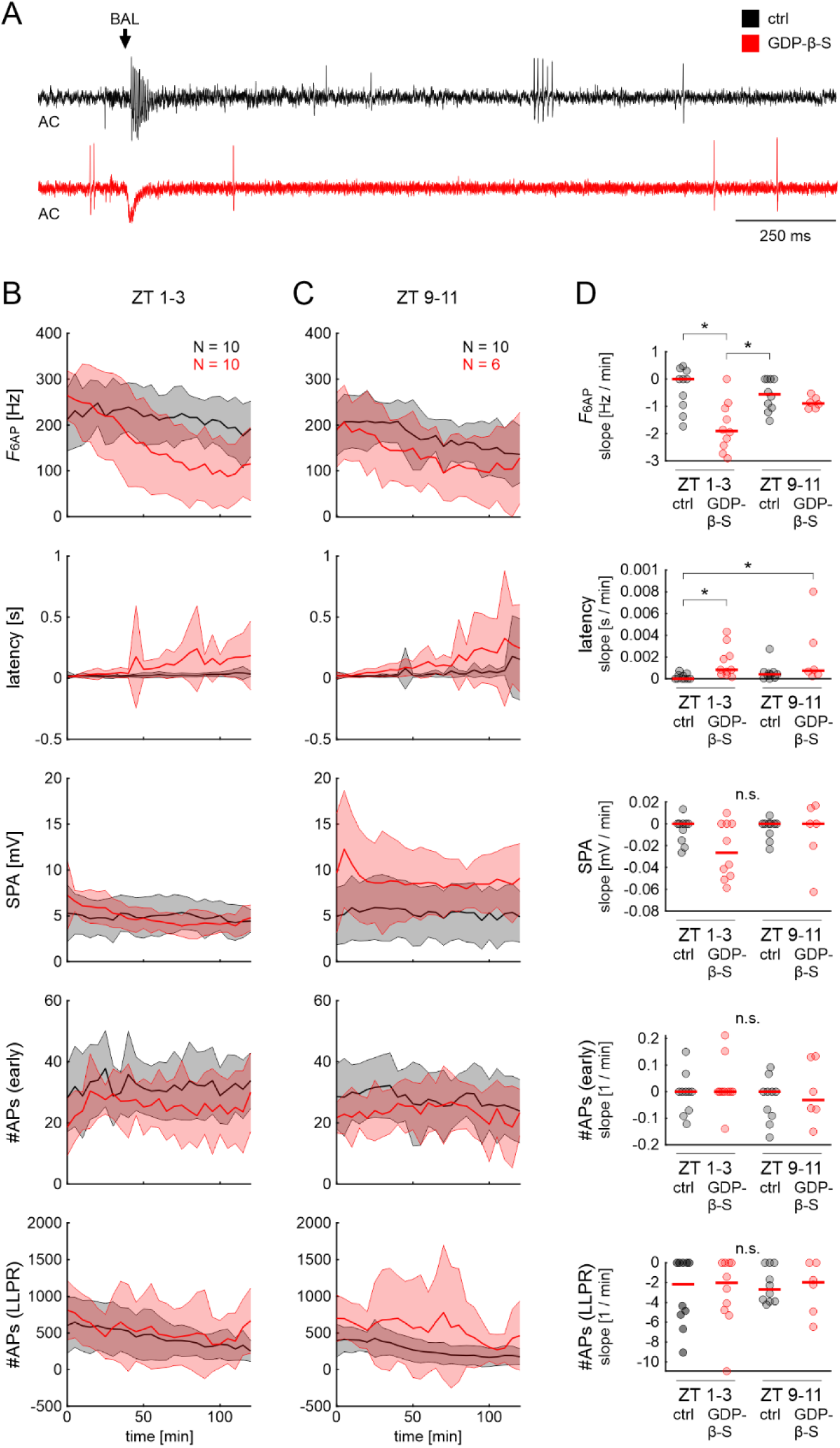
Inhibition of G protein signaling of pheromone sensitive ORNs by GDP-β-S decreased the phasic bombykal (BAL) response and increased response latency during the hawkmoth’s activity phase. **(A)** Examples of high-pass filtered tip-recordings (AC) of ORN responses in control (ctrl, recording solution + DMSO, black) and with the G protein antagonist GDP-β-S (dissolved in sensillum lymph ringer + DMSO, red) at ZT 1-3, 100 min after the start of the recording. Arrow indicates the onset of BAL response. Response parameters (see Figure 1) during the moth’s late activity phase (ZT 1-3; **B**) and at rest (ZT 9-11; **C**) in control and GDP-ϐ-S. Data are shown as mean (line) ± standard deviation (shaded area). **(D)** 1-way ANOVA results with appropriate post-hoc test for multiple comparisons (α = 0.05) for the slopes of BAL response parameters (see Methods and Figure 2A). *F*_6AP_ slopes decreased significantly, and latency slopes increased significantly in GDP-ϐ-S compared to control at the activity phase (ZT 1-3). No other parameters showed significant differences compared to control at either ZT. Dots show data for individual experiments, red lines indicate the mean. Raw data are provided in Extended Data Figure 3-1.

**Figure 4:**
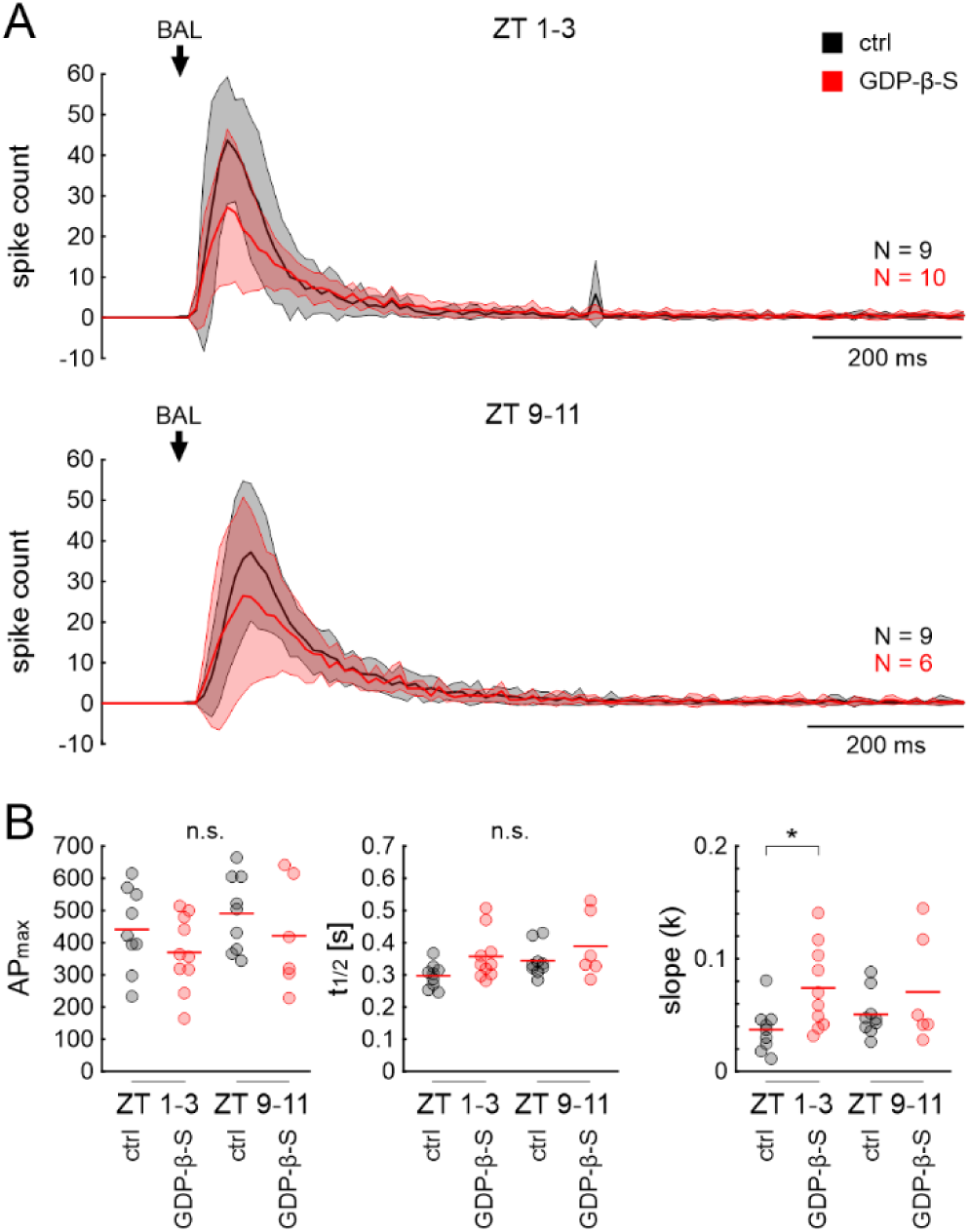
Inhibition of G protein signaling changed kinetics of the ORN response to BAL stimulation during the activity phase. **(A)** Peristimulus time histograms (PTSH; 10 ms bins) of the first second of the ORN response to BAL stimulation in control (black) and in GDP-ϐ-S (red). Data are shown as mean (line) ± standard deviation (shaded area). Arrow: onset of BAL response. **(B)** Fit parameters (total number of spikes in 1 s after onset of the BAL response (AP_max_), time of the sigmoid midpoint (t_1/2_) and slope (k) of the midpoint) of sigmoidal fits to the cumulative spike histograms (see Eq. 1, Methods, and Figure 2B). 1-way ANOVA with appropriate post-hoc test (α = 0.05) revealed a significantly steeper slope (k closer to zero) in control compared to GDP-ϐ-S during the activity phase (ZT 1-3). Steeper slopes indicate faster rise and fall times of the spike count in the PTSHs. The total number of spikes and sigmoid midpoint were not significantly different. Dots show fit parameter values for individual experiments, red lines indicate the mean. Raw data are provided in Extended Data Figure 4-1.

**Table 2:** 1-way ANOVA results for BAL response parameters in control and GDP-ϐ-S. If tests for normality (Shapiro-Wilk) or equal variance (Levene) failed, results are for ANOVA on ranks. The four groups are ctrl ZT 1-3, GDP-ϐ-S ZT 1-3, ctrl ZT 9-1, and GDP-ϐ-S ZT 9-11. p values smaller than α = 0.05 are printed in bold. N: number of animals; df: degrees of freedom; res: residual.

| N | parameter | normality | variance | test statistic | df | res. | p |
| --- | --- | --- | --- | --- | --- | --- | --- |
| 10 | $F_{6AP}$ | pass | pass | F = 7.492 | 3 | 32 | **<0.001** |
| 10 | latency | fail |  | H = 13.975 | 3 |  | **0.003** |
| 10 | SPA | pass | fail | H = 3.497 | 3 |  | 0.321 |
| 10 | #APs (early) | pass | pass | F = 0.546 | 3 | 32 | 0.654 |
| 10 | #APs (LLPR) | fail |  | H = 0.203 | 3 |  | 0.977 |

**Table 3:** 1-way ANOVA results for the fit parameters of the BAL response kinetics (see Eq. 1). Data for slope were log-transformed to ensure normal distribution. If tests for normality (Shapiro-Wilk) or equal variance (Levene) failed, results are for ANOVA on ranks. The four groups are ctrl ZT 1-3, GDP-ϐ-S ZT 1-3, ctrl ZT 9-1, and GDP-ϐ-S ZT 9-11. p values smaller than α = 0.05 are printed in bold. N: number of animals; df: degrees of freedom; res: residual.

| N | parameter | normality | variance | test statistic | df | res. | p |
| --- | --- | --- | --- | --- | --- | --- | --- |
| 10 | $AP_{max}$ | pass | pass | F = 1.414 | 3 | 30 | 0.258 |
| 10 | $t_{1/2}$ | fail | | H = 7.243 | 3 | | 0.065 |
| 10 | log(slope) | pass | pass | F = 3.212 | 3 | 30 | **0.037** |

In conclusion, only the first component of the BAL response, which encodes stimulus concentration (*F*_6AP_, latency) appears to be under the control of G proteins. This metabotropic signaling changed with ZT. The underlying differential effects that result in the increased variability of response parameters hint at more than one G protein coupled mechanism.

### 5.2 Inhibition of PLC reduced the spike frequency of the phasic component of the BAL response at both ZTs, while increasing response latency and decreasing SPA during the activity phase

In a complementary experiment, the PLC inhibitor U73122 was infused via the tip-recording electrode during the same ZTs as in the experiment above, where G protein action was blocked. Infusion of U73343, a structurally similar but very weak inhibitor of PLC, served as control. Overall, results were comparable to those obtained in GDP-ϐ-S with some ZT-dependent differences (Figure 5, ANOVA: Table 4; raw data: Extended Data Figure 5-1).

**Figure 5:**
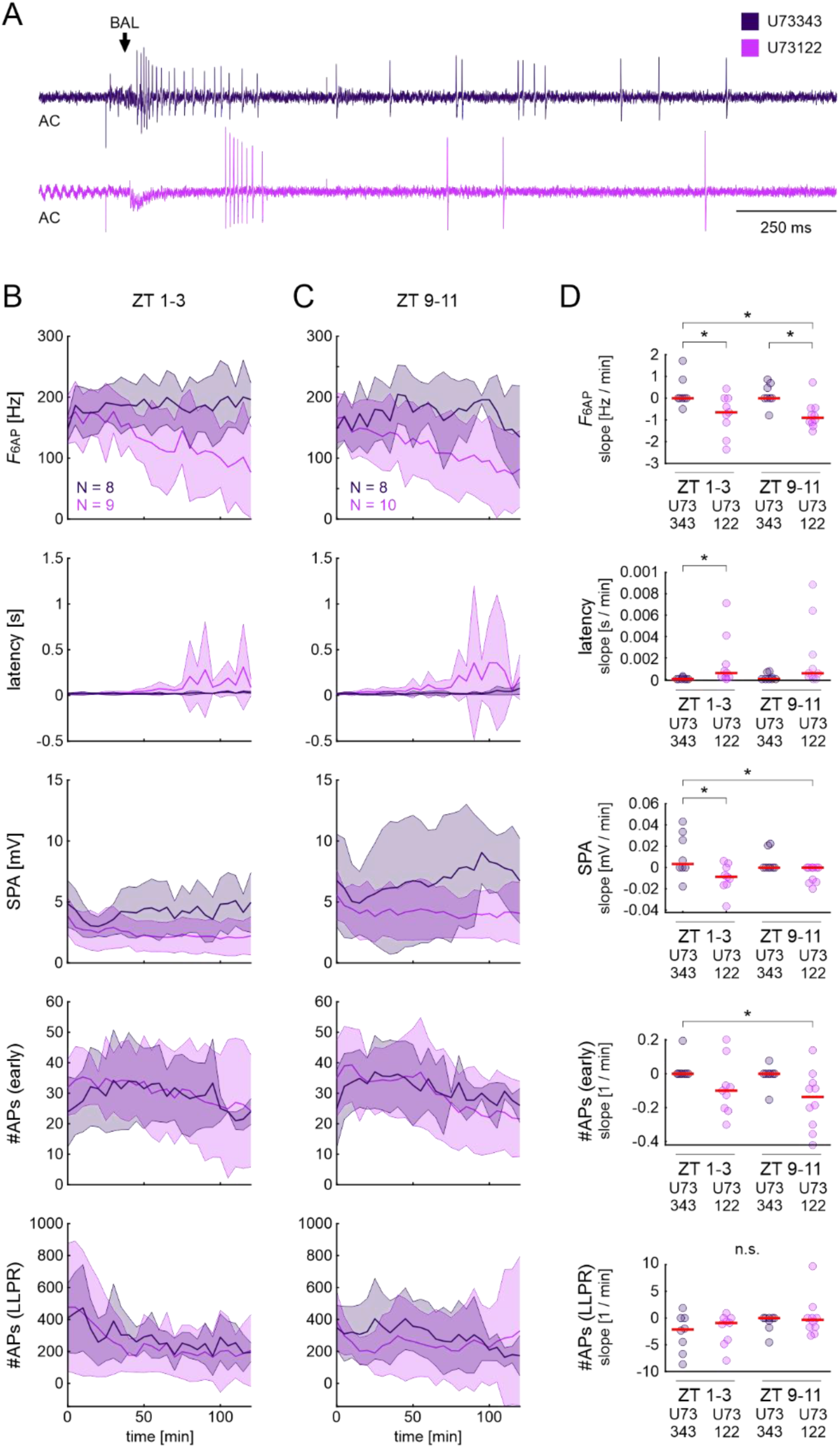
Inhibition of phospholipase C (PLC) of pheromone sensitive ORNs with U73122 decreased the phasic bombykal (BAL) responses at both ZTs, while decreasing sensillum potential amplitude (SPA) and increasing response latency only at ZT 1-3. **(A)** Example high-pass filtered (AC) tip-recordings of ORN responses in control (U73343, dark purple) and in PLC inhibitor (U73122, light purple) at ZT 1-3, 100 min. Arrow indicates the onset of the BAL response. Response parameters (see Figure 1) during the activity phase (ZT 1-3; **B**) and at rest (ZT 9-11; **C**) in control and U73122. Data are shown as mean (lines) ± standard deviation (shaded areas). **(D)** 1-way ANOVA results with appropriate post-hoc test for multiple comparisons (α = 0.05) for the slopes of BAL response parameters (see Methods and Figure 2A). Compared to controls, *F*_6AP_ slopes decreased at both ZTs in U73122 but SPA and latency slopes increased only at the activity phase (ZT 1-3). Other BAL response parameters showed no significant differences between control and U73122 at either ZT. Dots show data of individual experiments, red lines indicate the mean. Raw data are provided in Extended Data Figure 5-1

**Table 4:** 1-way ANOVA results for BAL response parameters in control (U73343) and U73122. If tests for normality (Shapiro-Wilk) or equal variance (Levene) failed, results are for ANOVA on ranks. Groups are ctrl ZT 1-3, U73122 ZT 1-3, ctrl ZT 9-11, and U73122 ZT 9-11. p values smaller than α = 0.05 are printed in bold. N: number of animals; df: degrees of freedom; res: residual.

| N | parameter | normality | variance | test statistic | df | res. | p |
| --- | --- | --- | --- | --- | --- | --- | --- |
| 10 | $F_{6AP}$ | pass | pass | $F = 5.447$ | 3 | 31 | **0.004** |
| 10 | latency | fail | | $H = 12.187$ | 3 | | **0.007** |
| 10 | SPA | pass | pass | $F = 4.493$ | 3 | 31 | **0.010** |
| 10 | #APs (early) | pass | fail | $H = 10.327$ | 3 | | **0.016** |
| 10 | #APs (LLPR) | pass | pass | $F = 1.558$ | 3 | 31 | 0.219 |

During the activity phase, the inhibition of PLC significantly reduced *F*_6AP_, increased latency, and reduced the SPA over time (Figure 5B, D). In addition, inhibition of PLC significantly reduced *F*_6AP_ over time during the resting phase (Figure 5C, D). All other response parameters did not show significant differences at either ZT. Similarly to the experiments with GDP-β-S, response variability across animals increased in U73122 at both ZTs over time, as indicated by the large increase in standard deviation (Figure 5B, C).

In conclusion, inhibition of PLC, a downstream target of G proteins, affected mainly the first component of the BAL response. These effects on PLC activation appeared to be under control of a circadian clock, since most of the effects were observed during the activity phase. Furthermore, differences in the effects between interference with general G protein signaling and specific PLC signaling hint at the involvement of more than one class of G protein in pheromone transduction (c.f. Figure 3, Figure 5).

### 5.3 Toxins targeting G_αo_ and G_αs_ reduced the spike frequency and increased the response latency of the phasic component of the BAL response during the activity phase

Thus far, the results indicated that G protein-signaling and activation of PLC were involved in the hawkmoth pheromone transduction cascade. Typically, PLC is activated by the αq subunit of G proteins and increases IP_3_ levels. Therefore, bacterial toxins were applied to specifically activate or inhibit different G_α_ subunits, which interferes with cAMP and IP_3_ levels. Cholera toxin (cTox) constitutively activates G_αs_, which activates adenylyl cyclase and leads to sustained increases in cAMP levels, also in insect antennae (Boekhoff et al., 1993; Deng et al., 2011). In vertebrates, Pertussis toxin (pTox) inhibits G_αo_ (Hopkins et al., 1988), a subunit that normally inhibits AC, which results in increases in cAMP levels (Mangmool and Kurose, 2011). In insect antennae, pTox has been shown to inhibit the pheromone-dependent IP_3_ production (Boekhoff et al., 1993, 1990). In vertebrates, *Pasteurella multocida* toxin (pmTox) activates both the G_αi/o_ subgroup and G_αq_, resulting in decreased cAMP and increased IP_3_ levels ((Surguy et al., 2014), not yet tested in insects). The following experiments were done only during the hawkmoth’s activity phase at ZT 1-3 because G protein and PLC inhibition in the previous experiments were effective at those times. As with the chemicals before, toxins were applied to the sensillum via the recording electrode.

Both cTox and pTox application resulted in significant differences in the first and second component of the BAL response that were similar to the results obtained with GDP-β-S and U73122: Activation of G_αs_ and inhibition of G_αo_ both decreased *F*_6AP_ and increased latency significantly (Figure 6; ANOVA: Table 5; raw data: Extended Data Figure 6-1). In addition, these toxins significantly reduced the number of spikes in the first and second component of the response (#APs (early)). SPA and #APs (LLPR) were not altered. The application of pmTox did not have any significant effects (Figure 6B).

**Figure 6:**
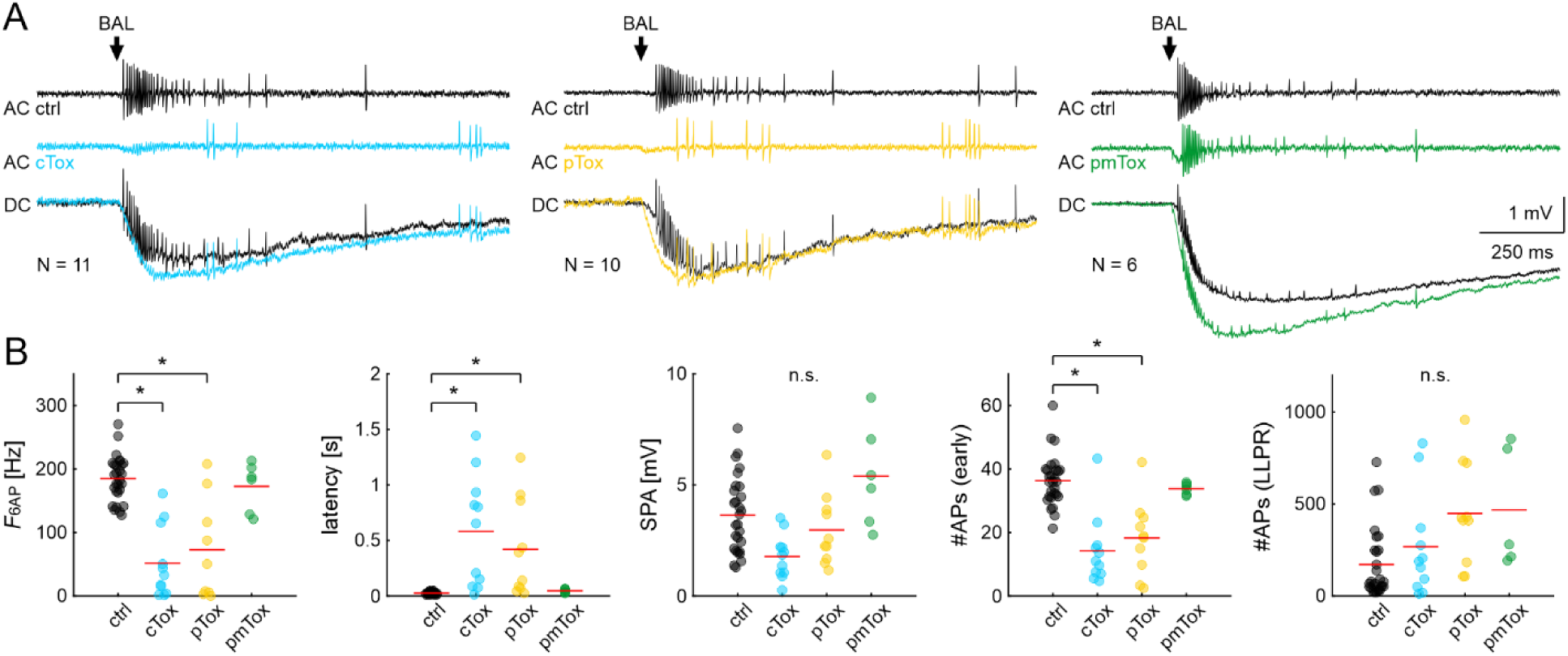
Responses to bombykal (BAL) stimulation at ZT 1-3 were affected by toxins that target Gα_s_ or Gα_o_ subunits. **(A)** Examples of tip-recordings from ORNs with BAL stimulation in control (black), with cholera toxin (cTox; blue), pertussis toxin (pTox; yellow), and *Pasteurella multocida* toxin (pmTox, green); repeated measures of one animal shown for each toxin. AC: high-pass filtered recording to highlight action potential (AP) response; DC: unfiltered sensillum potential amplitude (SPA) with APs. Arrow: onset of the BAL response. **(B)** Quantification with 1-way RM ANOVA with appropriate post-hoc test for multiple comparisons (α = 0.05) of the same parameters as in Figure 3 and Figure 5. During the activity phase (ZT 1-3), the frequency of the phasic BAL responses (F_6AP_) and the number of Aps during the first 100 ms of the response (#APs (early)) decreased in cTox (blue; sustained activation of Gα_s_) and pTox (yellow; inhibition of Gα_o_), while response latency increased significantly. The effect of pmTox (green; constitutive activation of G_α12/13_, G_αi_, and G_αq_) was not different from control. Raw data are provided in Extended Data Figure 6-1.

**Table 5:** 1-way RM ANOVA results for BAL response parameters in control and bacterial toxins at ZT 1-3. Values for latency were log-transformed to ensure equal variance. Groups are ctrl, pTox, cTox, pmTox. p values smaller than α = 0.05 are printed in bold. N: number of animals; df: degrees of freedom; res: residual.

| N | parameter | normality | variance | test statistic | df | res. | p |
| --- | --- | --- | --- | --- | --- | --- | --- |
| 28 | $F_{6AP}$ | pass | pass | F = 26.611 | 3 | 23 | **<0.001** |
| 28 | log(latency) | pass | pass | F = 26.606 | 3 | 25 | **<0.001** |
| 28 | SPA | pass | pass | F = 1.996 | 3 | 24 | 0.142 |
| 28 | #APs (early) | pass | pass | F = 22.252 | 3 | 24 | **<0.001** |
| 28 | #APs (LLPR) | pass | pass | F = 3.726 | 3 | 23 | **0.026** |

In conclusion, G protein-dependent activation/disinhibition of adenylyl cyclase affected the BAL transduction in hawkmoths in the same way as PLC inhibition. Since pmTox did not result in significant changes of the BAL response, it remains to be studied whether pmTox affects any G proteins in hawkmoth. The application of the toxins that increase cAMP concentration replicated the results that we obtained from G protein and PLC inhibition, indicating that G protein activation and cyclic nucleotides are involved in the regulation of ORN sensitivity to pheromones.

### 5.4 Expression levels of PLCϐ4 peaked during the activity phase

During the activity phase of the hawkmoth, but not during the resting phase, the disruption of G proteins and PLC affected the pheromone response parameters that encode BAL concentration. Therefore, we examined whether these effects were caused by circadian changes in the antennal expression levels of G_αo_ or G_αq_, which activate PLCβ (increased IP_3_ levels), or G_αs_, which activate adenylyl cyclase (increased cAMP levels). The expression levels of G_α_ subunits and PLCβ, which we found in the transcriptome of male hawkmoth antennae, were quantified with qPCR (Figure 7).

**Figure 7:**
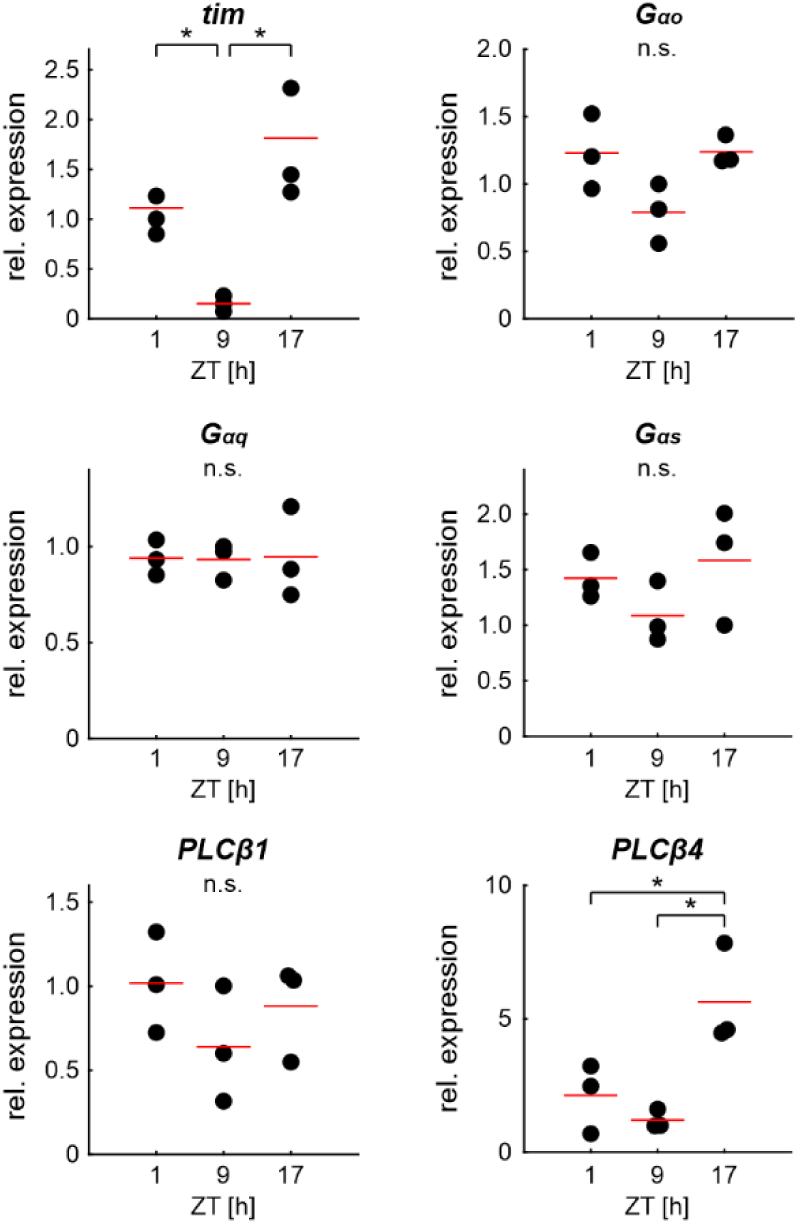
Relative expression levels of mRNA for G proteins, PLCβ, and the circadian clock protein timeless (*tim*) in male hawkmoth antenna at different zeitgeber times (ZTs). qPCR revealed that mRNA levels of *PLCϐ4* peaked significantly at the beginning of the activity phase at ZT 17. G_αo_, G_αq_, G_αs_, and *PLCϐ1* did not change throughout the day. The mRNA of the cycling circadian clock protein timeless (*tim*) served as positive control. Dots indicate values for biological replicates, each biological replicate contains extracts from 8 antennae and three technical repeats. Red lines depict the mean. 1-way ANOVA with appropriate post-hoc test for pairwise comparisons (α = 0.05). The phylogenetic tree for PLCβ is provided in Extended Data Figure 7-1. The nucleotide sequences of the genes are listed in Extended Data Figure 7-2.

Phylogenetic analysis clustered the putative PLCβ candidate transcripts with those of *B. mori* (Extended Data Figure 7-1; the nucleotide sequences of the genes are listed in Extended Data Figure 7-2). Expression of the circadian clock gene *timeless* (*tim*) served as positive control. Only *tim* and PLCβ4 (*norpA*) showed significantly different expression levels throughout the day, with higher expression levels during the activity phase (Figure 7; Table 6). The expression of G_α_ subunits and PLCβ1 did not change.

**Table 6:**
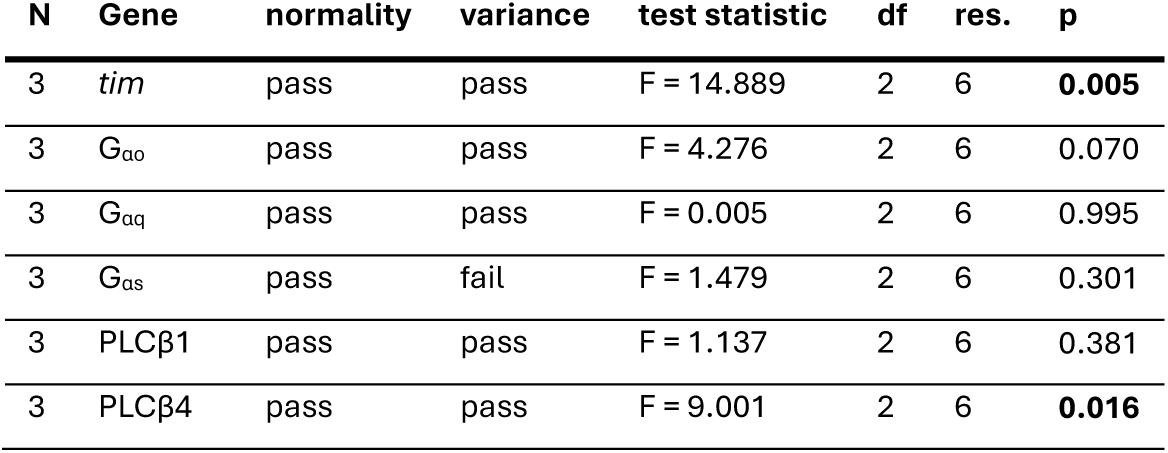
1-way ANOVA results for gene expression levels at ZT 1, ZT 9, and ZT 17. p values smaller than α = 0.05 are printed in bold. N: number of animals; df: degrees of freedom; res: residual.

| N | Gene | normality | variance | test statistic | df | res. | p |
| --- | --- | --- | --- | --- | --- | --- | --- |
| 3 | *tim* | pass | pass | F = 14.889 | 2 | 6 | **0.005** |
| 3 | $G_{ao}$ | pass | pass | F = 4.276 | 2 | 6 | 0.070 |
| 3 | $G_{aq}$ | pass | pass | F = 0.005 | 2 | 6 | 0.995 |
| 3 | $G_{as}$ | pass | fail | F = 1.479 | 2 | 6 | 0.301 |
| 3 | PLCβ1 | pass | pass | F = 1.137 | 2 | 6 | 0.381 |
| 3 | PLCβ4 | pass | pass | F = 9.001 | 2 | 6 | **0.016** |

## 6 Discussion

Chemosensory transduction is under strict circadian control in insects, and previous experiments in solitary nocturnal moths suggest that the transduction cascade differs between sleep and activity states (Dolzer et al., 2021; Flecke et al., 2010, 2006; Gawalek and Stengl, 2018; Krishnan et al., 1999; Linn et al., 1996; Merlin et al., 2007; Page and Koelling, 2003; Plautz et al., 1997; Rosén et al., 2003; Rymer et al., 2007; Zhukovskaya, 1995) In *D. melanogaster* ORNs, experimental data support an ionotropic odor transduction cascade in addition to cAMP-dependent metabotropic signaling in response to general odorants (Nakagawa and Vosshall, 2009; Wicher and Miazzi, 2021). In contrast, no evidence was found for OR-Orco-dependent ionotropic pheromone transduction in the hawkmoth *M. sexta*, but evidence accumulated for G protein-coupled activation of phospholipase Cβ (PLCβ) (Nolte et al., 2016, 2013; Stengl, 2010; Stengl and Funk, 2013).

Although the inverse topology of insect ORs raised doubts about potential G protein coupling (Benton et al., 2006; Wistrand et al., 2006), and some physiological studies found no evidence for G protein-dependent cascades (Sato et al., 2008; Yao and Carlson, 2010), several physiological and neuroanatomical findings in different insect species strongly suggest that metabotropic cascades are involved in insect odor/pheromone transduction (Boekhoff et al., 1993, 1990; Boto et al., 2010; Breer et al., 1990, 1988; Chatterjee et al., 2009; Deng et al., 2011; Fleischer et al., 2018; Grosse-Wilde et al., 2010; Ignatious Raja et al., 2014; Kain et al., 2008; Kalidas and Smith, 2002; Laue et al., 1997; Nakagawa and Vosshall, 2009; Stengl, 2010, 1994, 1993; Stengl and Funk, 2013; Wegener et al., 1997; Wetzel et al., 2001; Wicher et al., 2008; Wicher and Miazzi, 2021; Ziegelberger et al., 1990). Therefore, in this study, we combined pharmacological experiments, which disrupted metabotropic signaling through different subunits of G_α_ proteins and their target, PLCβ, with tip-recordings of pheromone-sensitive trichoid sensilla in intact, healthy hawkmoths. Our qPRC results provide evidence for circadian control of BAL-dependent transduction cascades.

### 6.1 The highly sensitive pheromone transduction in moths is coupled to G_αo_ and PLCϐ

Solitary nocturnal moth males have only a few weeks as adults to accomplish the challenging task of locating their species-specific females for mating. Thus, evolution equipped moths with an exquisitely sensitive sense of smell that apparently allows the detection of single pheromone molecules (Kaissling, 1986; Kaissling and Priesner, 1970). To do so, the system needs mechanisms for the reduction of noise in the form of other olfactory molecules and maximized odor-specific signal amplification. Patch clamp experiments in primary cell cultures of *M. sexta* ORNs revealed the activation of three consecutive pheromone-dependent ionic currents with physiologically low concentrations of pheromone stimuli (Stengl, 1994, 1993). First, a rapid, transient, pheromone-dependent Ca^2+^ inward current activates and declines in less than 100 ms after the start of the pheromone response. Second, apparently based on the resulting increase in intracellular Ca^2+^, a slower Ca^2+^-gated, non-specific cation current activates, which underlies the slower kinetics of the second component of the pheromone response. Third, a protein kinase C -dependent, slow, non-specific cation current activates that persists for several second and underlies the third component of the pheromone response (Stengl, 2010). Since that first Ca^2+^ channel of the transduction cascade functions as a “yes-or-no trigger” for subsequent Ca^2+^-dependent amplification steps, it both obliterates noise of previous processes and amplifies the signal via Ca^2+^-dependent mechanisms.

The properties of that first Ca^2+^-channel are different from those of the OR-Orco cation channels of *D. melanogaster* (Sato et al., 2008; Wicher et al., 2008), so that OR-Orco-dependent ionotropic signaling seems unlikely in hawkmoth pheromone transduction (Nolte et al., 2016, 2013). Rather, the channels that give rise to at least the first two of the three consecutive currents of the pheromone response in *M. sexta* seem to belong to the TRPC superfamily of TRP channels that are highly permeable to Ca^2+^, which were originally described in the similarly highly sensitive phototransduction cascade of *D. melanogaster* (Gawalek and Stengl, 2018; Hardie and Minke, 1992). It was discovered only recently that TRPCs are not only linked to PLCϐ activation but can also be directly gated by G_i/o_ proteins (Jeon et al., 2020; Zhu, 2023). Whether this is true for the antennal TRPs of *M. sexta* remains to be examined. Only the late, long-lasting third component of the pheromone response was linked to Orco in *M. sexta*. This makes an ionotropic mechanism of pheromone transduction unlikely and rather suggests cGMP-dependent and/or voltage-dependent Orco gating (Nolte et al., 2016, 2013).

The findings described in the current study are consistent with localization and functional analyses of G protein subunits and PLCβ in different insect antennae. In antennae of *D. melanogaster*, RT-PCR reported the expression of six G_α_ subunits (G_s_, G_i_, G_q_, G_o_, G_f_, and concertina) in addition to different G_ϐ_ and G_γ_ subunits (Boto et al., 2010). There, G_s_, G_i_, G_q_ are located in the cilia of ORNs, suggesting coupling to ORs (Boto et al., 2010; Kain et al., 2008). Different physiological and behavioral assays, in combination with molecular genetics, demonstrated that G_αo_ or G_αq_ signaling is used in odor detection in *D. melanogaster* (Chatterjee et al., 2009; Ignatious Raja et al., 2014; Kain et al., 2008; Kalidas and Smith, 2002). In contrast, Yao and Carlson (2010) reported in an extensive study that G_αq_ and G_γ30A_ are involved in CO_2_ detection through gustatory receptors, but not in OR-mediated odor signaling of antennal basiconic sensilla.

Regarding moths, G_αo_ G_αq_, and G_αs_ are expressed and localized in adult antennae of *B. mori* (Miura et al., 2005). In *Antheraea pernyi*, G_αq_ was found in homogenates of the antennal ciliary fraction and, by electron-microscopic studies, was detected in cilia of pheromone-sensitive trichoid sensilla. Hence, it is likely that G_αq_ is involved in pheromone detection in *A. pernyi* by coupling to ORs (Laue et al., 1997).

Furthermore, the pheromone-dependent fast, transient rises of IP_3_ in homogenates of moth and cockroach antennae are consistent with the pheromone-dependent activation of PLCϐ during the first component of the pheromone response (Boekhoff et al., 1993; Breer et al., 1990). Since pheromone-dependent rises in IP_3_ levels are decreased by interfering with G_αo_ through pTox in the antennae of insects (Boekhoff et al., 1990), it is likely that, consistent with current data in hawkmoths and our results presented here, pheromone receptors in moths use G_αo_-dependent activation of PLCβ. It remains to be tested in detail whether and how pheromone transduction of different insect ORs couple directly to G proteins and their respective target, PLCβ, or to other enzymes such as adenylyl cyclases.

In *D. melanogaster* and other insect species, all members of the cAMP signaling cascades have been found in the antennae, located in ORNs, by either demonstrating the respective gene expression with molecular genetic tools, or by morphological assays with immunocytochemistry. Rises in cAMP were reported after odor stimulation and mutations of the cAMP cascade results in defects in olfaction-dependent behavior, which suggests that at least some specific ORs are relying on G_αs_-dependent activation of adenylyl cyclases in *D. melanogaster* (Deng et al., 2011; Wicher et al., 2008; Wicher and Miazzi, 2021).

The discrepancies in reports of insect olfaction could stem from inconsistent experimental conditions between the different studies. Conclusions from measurements of different sensillum types and ORNs are often generalized and not distinguished in different or even the same species. Additionally, odorant concentrations and stimulus durations differ vastly between publications.

Furthermore, the intracellular Ca^2+^ concentrations in the recorded ORNs are not controlled and would affect ion channel availability for odor-dependent activation (Stengl, 2010; Stengl and Funk, 2013). As we demonstrated here, interference with pheromone transduction cascades depended on ZT, and mostly affected the phasic component of the pheromone response. This and previous studies indicate that the three components of the response are mediated by distinct mechanisms of the transduction cascade, and failure to address these components separately could obscure G protein actions that seem to act mostly on the first component. Very likely, careful planning and comparative analysis of experiments across species will show that there are several redundant transduction mechanisms for insect ORs, depending on the different physiological needs of the respective insect.

In summary, our physiological data from *in vivo* recordings of hawkmoth trichoid sensilla are consistent with pheromone receptor-dependent activation of G_αo_, which causes activation of PLCβ4, and therefore increases of IP_3_ and diacylglycerol levels. Although BAL receptors in hawkmoth have the same inverse 7TM topology as OR in other insects (Grosse-Wilde et al., 2007, 2006; Wicher et al., 2017), our results indicate direct or indirect coupling to G protein-dependent amplification cascades, as shown for insect gustatory receptors and ionotropic receptors (Ishimoto et al., 2005; Nakagawa et al., 2005; Stengl, 2010; Ueno et al., 2006; Wetzel et al., 2001; Yao and Carlson, 2010). It seems that, in hawkmoths, there are several redundant or interdependent transduction cascades that guarantee the reliable, highly sensitive pheromone transduction, which depends on odor stimulus strength, duration, time course, and time of day, as well as on intracellular second messengers that vary across the day (Stengl, 2010; Stengl and Funk, 2013; Stengl and Schröder, 2021). It remains to be studied if and how receptors in ORNs and even in the non-neuronal support cells interact during pheromone and odor transduction.

### 6.2 Peripheral circadian clocks orchestrate olfaction-dependent behavioral rhythms via circadian control of chemosensory responsiveness

Nocturnal male moths are maximally responsive to pheromones during their activity phase in the scotophase (Flecke et al., 2010; Linn et al., 1996; Rosén et al., 2003). These rhythms in responsiveness are controlled by circadian clocks and therefore persist under constant conditions (Baker and Cardé, 1979; Merlin et al., 2007; Rosén et al., 2003). Similarly, females, guided by their own circadian clocks, release pheromones at night, and peak release correlates with the males’ search flight (Itagaki and Conner, 1988; Rosén, 2002; Sasaki and Riddiford, 1984). Mating success decreases if males and females are raised out of phase in different light-dark cycles (Silvegren et al., 2005). Both photoperiod and pheromone presence synchronize the mating behavior of moths by circadian modulation of female pheromone release and male pheromone sensitivity. How and where exactly this circadian modulation occurs in the male’s olfactory system remains to be investigated further. Our results indicate that PLCβ4 is under circadian control and could be engaged in the heightened pheromone sensitivity, consistent with previous finding of circadian changes of IP_3_ levels in hawkmoth antennae (Schendzielorz et al., 2015).

Circadian clocks generate rhythms with a cycle period of approximately 24 hours through positive feedforward and delayed negative feedback loops. These clocks are formed at the gene transcription level as transcriptional/translational feedback loop (TTFL) oscillators, or at the posttranslational level as posttranslational feedback loop (PTFL) oscillators (Stengl and Schneider, 2024). The widespread and intertwined TTFL and PTFL oscillators sustain physiological homeostasis (Stengl and Schneider, 2024) and occur not only in the central and peripheral nervous system but also in many organs throughout an organism (Giebultowicz, 2000; Plautz et al., 1997).

In response to food odors and pheromones, electroantennogram (EAG) recordings from *D. melanogaster* and the cockroach *Rhyparobia maderae* show rhythmic activity (Krishnan et al., 1999; Merlin et al., 2007; Page and Koelling, 2003; Rymer et al., 2007). Only targeted ablation of TTFL clock genes in the antenna, but not ablation of brain clocks, did affect the circadian rhythms of EAGs in *D melanogaster*, showing that the circadian oscillators in the antennae are necessary and sufficient for chemosensory rhythms (Tanoue et al., 2004). Which mechanisms and which antennal cells control circadian rhythms in odor and pheromone sensitivity in insects? Besides the ORNs in *M. sexta*, non-neuronal supporting cells and epithelial cells in the antennal sensilla express circadian clock genes with circadian rhythms, which suggests that many intertwined clocks and processes govern pheromone sensitivity in hawkmoths (Schuckel et al., 2007). Antennal genes that are involved in odorant binding and odorant clearance are rhythmically expressed (Ceriani et al., 2002; Claridge-Chang et al., 2001; Dhungana et al., 2023; Jin et al., 2017; McDonald and Rosbash, 2001; Merlin et al., 2007; Saifullah and Page, 2009). We could show for *M. sexta* that PLCβ4 expression depends on the time of day, which makes it a candidate for the circadian control of chemosensory sensitivity.

## Acknowledgements

We thank Niklas Metzendorf and Jan Bröckel for helpful discussions of this manuscript, and James H. Tumlinson for his generous gift of synthetic bombykal. ChatGPT has been used for spell-checking in the qPCR related sections of Methods.

## 9 Extended Data Legends

**Extended Data Figure 3-1**

All data for the analysis shown in Figure 3. The excel file contains five sheets: one with the slopes and four with the raw data for the four combinations of ZT and experimental conditions from which the slopes were fitted. “slopes” contains the two datasets at ZT 1-3 and ZT 9-11. In each dataset, the first column is the unique experiment ID, the second column denotes the experimental condition, and the following columns give the slope for the five response parameters. In the other four sheets, the first column denotes the stimulation time in minutes after the electrodes were attached. Here, data are organized by experiment ID and show the value of the respective response parameter for each stimulation time point. “NaN” indicates those trials where we could not extract the relevant parameters from the recording.

**Extended Data Figure 4-1**

All data for the analysis shown in Figure 4. Each dataset is a combination of ZT and experimental condition. For each dataset, the first column denotes the experiment ID, and the following three columns the respective fit parameters.

**Extended Data Figure 5-1**

All data for the analysis shown in Figure 5. The excel file contains five sheets: one with the slopes and four with the raw data for the four combinations of ZT and experimental conditions from which the slopes were fitted. “slopes” contains the two datasets at ZT 1-3 and ZT 9-11. In each dataset, the first column is the unique experiment ID, the second column denotes the experimental condition, and the following columns give the slope for the five response parameters. In the other four sheets, the first column denotes the stimulation time in minutes after the electrodes were attached. Here, data are organized by experiment ID and show the value of the respective response parameter for each stimulation time point. “NaN” indicates those trials where we could not extract the relevant parameters from the recording.

**Extended Data Figure 6-1**

All data for the analysis shown in Figure 6. The excel file contains three sheets, one for each toxin condition. Data are grouped by experiment ID. The first column contains the stimulation time points. Subsequent columns contain the respective response parameter at that time point. Experiments were paired with control recordings on day 1 and toxin recordings on day 2. “NaN” indicates those trials where we could not extract the relevant parameters from the recording.

**Extended Data Figure 7-1**

The phylogenetic tree was constructed using the maximum likelihood method, and node support was evaluated with 1000 bootstrap replicates. Bootstrap values > 67 are shown. Black boxes and black circles represent candidate PLCβ1 and PLCβ4 of *M. sexta* (Gene IDs: LOC115440592 and LOC115451385), respectively. Sequence information is detailed in Extended Data Figure 7-2.

**Extended Data Figure 7-2**

Nucleotide sequences for the genes used in the phylogenetic analysis.

