## Supplementary figures and images for "Hawkmoth pheromone transduction involves G protein-dependent phospholipase Cβ signaling"

### Extended Data Figure 7-1

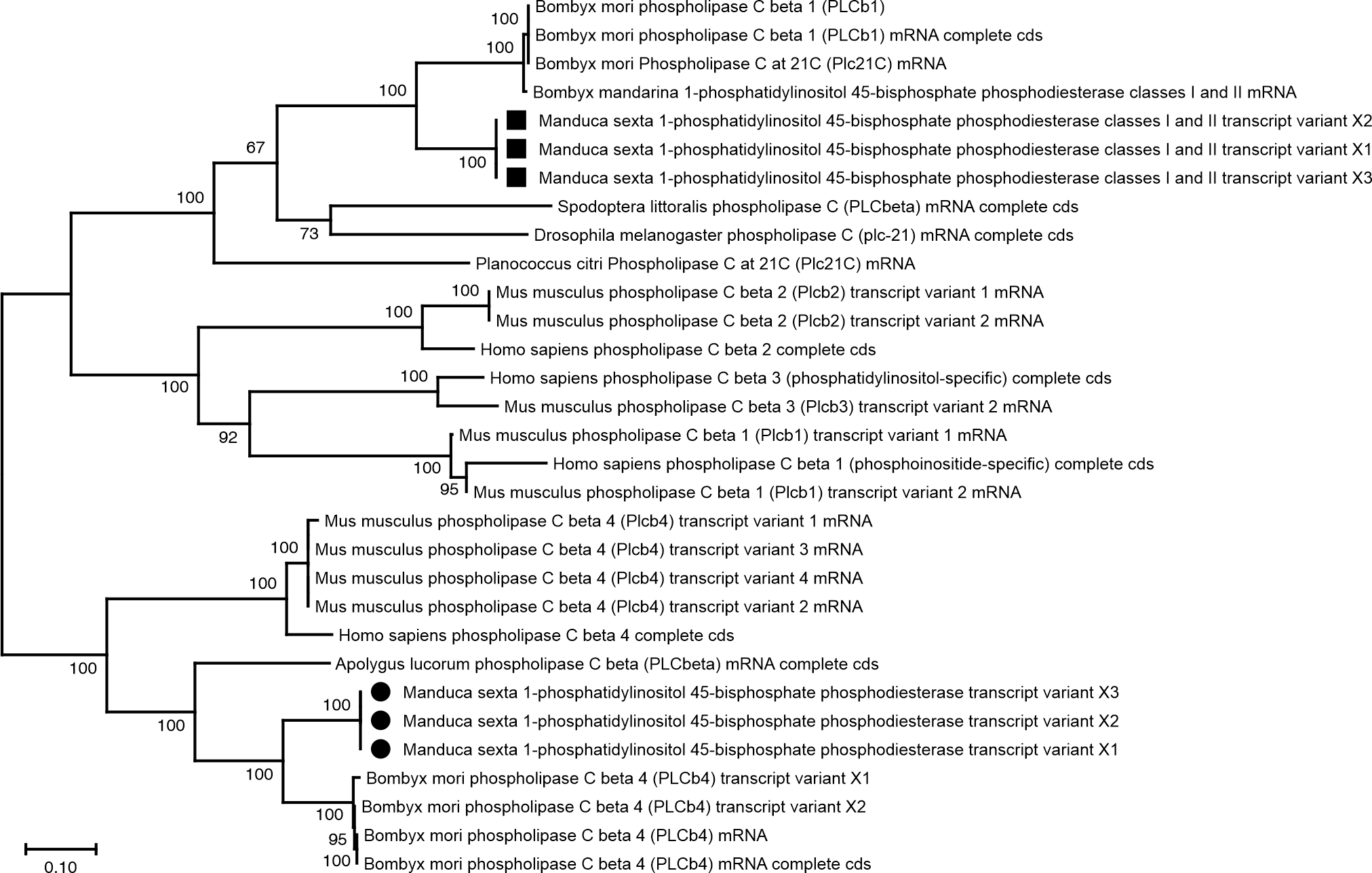
